# A conserved genetic basis for commensal-host specificity through live imaging of colonization dynamics

**DOI:** 10.1101/2024.04.19.590229

**Authors:** Karina Gutiérrez-García, Kevin Aumiller, Ren Dodge, Benjamin Obadia, Ann Deng, Sneha Agrawal, Xincheng Yuan, Richard Wolff, Nandita Garud, William B Ludington

## Abstract

Animals throughout the metazoa selectively acquire specific symbiotic gut bacteria from their environment that aid host fitness. Current models of colonization suggest these bacteria use weakly specific receptors to stick to host tissues and that colonization results when they stick in a region of the host gut that overlaps with their nutritional niche. An alternative model is that unique receptor-ligand binding interactions provide specificity for target niches. Here we use live imaging of individual symbiotic bacterial cells colonizing the gut of living *Drosophila melanogaster* to show that *Lactiplantibacillus plantarum* specifically recognizes a distinct physical niche in the host gut. We find that recognition is controlled by a colonization island that is widely conserved in commensals and pathogens from the Lactobacillales to the Clostridia. Our findings indicate a genetic mechanism of host specificity that is broadly conserved.

**One-Sentence Summary:** Host-symbiont specificity is encoded by a conserved colonization island that provides molecular precision to host niche access.

## Main Text

Bacterial colonization of the animal gut assembles a microbiome that aids in digestion (*1, 2*), vitamin production (*3*), host immune modulation (*4*), and prevention of pathogen invasion (*5*). Colonization depends in part on the physiological demands of the bacteria, including nutrients, pH and O_2_ requirements, their resistance to host factors, including stomach acid, bile salts, and immune effectors, as well as successful competition with other bacteria strains (*6*). The combination of all these host and bacterial factors results in “ecological filtering,” where the set of colonized bacteria are those that entered the gut and persisted (*5, 7*).

Along the length of the animal gut, there is considerable physiological variation in support of distinct host digestive processes such as protein decomposition and lipid absorption. Correspondingly, there is considerable spatial variation in bacterial composition along the digestive tract, with regions such as the mouth, stomach, jejunum, and colon having distinct compositions (*8*–*10*). Hosts can control the localization of bacteria by providing a niche site that sequesters and maintains specific bacteria (*11*–*14*). Bacteria also use proteins called adhesins, which stick to host tissues and prevent washout by peristaltic flow (*15*). Adhesins are operationally defined and do not have a conserved structure; they often bind somewhat non-specifically, based on electrostatic interactions (*15*). While specific binding of terminal mannose glycosylations in the urinary tract is known from an Enterobacterial pathogen (*16*), opportunistic pathogens are not selected over long evolutionary time for their ability to cause infections (*17*), and conserved mechanisms of host specificity are not known to regulate symbiont colonization. However, cophylogenetic patterns of evolution between symbionts and their hosts (*18*–*20*) suggest these mechanisms exist. How hosts and symbionts recognize one another to maintain site-specific colonization remains poorly understood even in the best studied models.

Genomic islands that promote host association are best studied in opportunistic pathogens, including *Streptococcus* and *Staphylococcus*, which are lactic acid bacteria related to the lactobacilli (*21*–*23*). Symbiosis islands are also known from symbiotic bacteria (*24*) including the *Vibrio fischeri* strains that colonize the light organs of squid (*25, 26*). These islands carry clusters of genes that are likely involved in bacterial association with their host and often contain adhesins (*27*).

Differentiating transient binding due to non-specific stickiness, as is often observed in opportunistic pathogens, from specific adhesion by a symbiont to niche tissue is complicated because it occurs deep inside the host gut, where other selective pressures such as nutrient availability act simultaneously. Live imaging studies, for instance in zebrafish larvae (*28*), can directly observe bacterial cells as they migrate through the gut (*29*). We developed single bacterial cell tracking in living *Drosophila* to differentiate between static adhesion to a precise physical location in the gut versus transient adhesion. *Drosophila melanogaster* fruit flies are an emerging model system for the gut microbiome, with a relatively low diversity of 5 to 20 species, including Firmicutes and Proteobacteria (*30*). *Lactiplantibacillus plantarum*, one of the widest-used human and animal probiotics (*31, 32*), is one of the most consistently associated fly bacteria across many studies (*33*–*35*), and it colonizes a precise, spatially-defined physical niche in the foregut of the fly (*12*). Despite over a century of work on *L. plantarum*, a host specific adhesion mechanism has never been identified, and *L. plantarum* has been labeled nomadic (*36*). Here, we investigated the physical and genetic basis for fly gut colonization specificity by *L. plantarum* and found broad conservation of the necessary genes across the Firmicutes, one of the most-host-associated bacterial phyla but for which host specificity genes are almost completely unknown.

### *Lactobacillus* colonizes by stably binding the niche

*L. plantarum WF* (*LpWF*), isolated from the gut of a wild *D. melanogaster* colonizes the gut with higher efficiency than related strains of *L. plantarum* isolated from humans, lab flies, and silage (**Fig. 1A-F, Fig. S1**; (*12, 37*)). To investigate the stability of colonization at the single bacterial cell level, we directly visualized individual mCherry-expressing bacterial cells colonizing the niche in living flies by optimizing the Bellymount video microscopy technique (*38*) at a focal plane in close proximity to the inner surface of the crop (**Fig. 1G-H, Fig. S2**). As a control, we imaged the weak colonizer, *LpATCC8014* (**Fig. 1I**; **Movie S1**), which diffused with a median diffusion coefficient of 0.10 µm^2^/s (**Fig. 1J**), consistent with the theoretical prediction from the Stokes-Einstein equation (**Fig. S2**). *L. plantarum* lacks cellular machinery for motility, such as flagella (*39*), so diffusive movement is expected. In *LpATCC8014*, cases of transient binding were observed followed by detachment, consistent with the observed probabilistic colonization (*37*). In contrast, *LpWF*, cells adhered to the inner surface of the crop (**Fig. 1H, K, Movie S2**) with a median diffusion coefficient of 0.001 µm^2^/s. We observed no detachment events for *LpWF*. Thus, *LpWF* cells colonize by static adhesion rather than transiently adhering and rebinding. We note that this binding occurs only in the symbiotic niche tissue and not in any other part of the gut, including the midgut (*12*).

**Figure 1.**
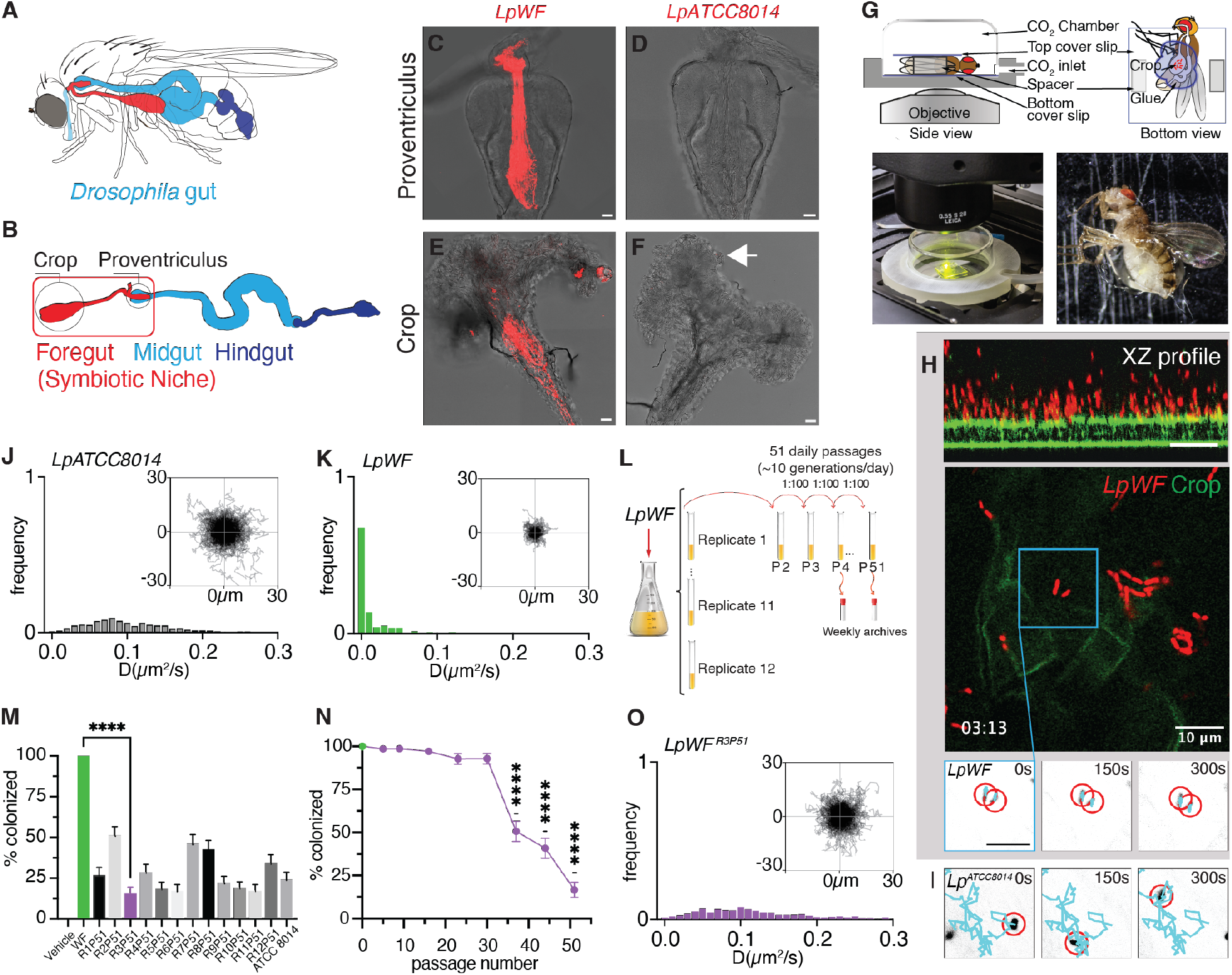
Live imaging of bacterial cells in the Drosophila gut shows that *Lactiplantibacillus plantarum WF* (*LpWF*) attaches specifically to a physical niche in the host foregut. (**A, B**) Schematics showing the anatomy of the fruit fly gut in the fly (A) and dissected (B), with the symbiotic niche highlighted in red. (**C**) Confocal micrograph of a proventriculus colonized by mCherry-labeled *LpWF*. (**D**) Proventriculus colonized by mCherry-labeled *LpATCC8014*. (**E**) Crop colonized by mCherry-labeled *LpWF*. (**F**) Crop colonized by mCherry-labeled *LpATCC8014*. Red: mCherry fluorescence. Note that *LpWF* is a colonizer and *LpATCC8014* is not. (**G**) Schematic and photos showing the Bellymount apparatus for live imaging. (**H**) X-Z (top), X-Y (middle) and X-Y-time-series (bottom) images of *LpWF* by Bellymount. (**I**) X-Y-time-series and particle tracking of *LpATCC8014*. (**J**) Histogram of coefficients of diffusion for individual *LpATCC8014* and (**K**) *LpWF* cells, n=500 tracks each from 5 biological replicates. (**L**) Schematic describing the experimental evolution screen. (**M**) Percentage of flies colonized by replicate 3 evolved mutants over time, n ≥ 72 individual flies from 3 biological replicates per strain. (**N**) Percent of flies colonized by evolved strains diminishes with passage number, n ≥ 72 individual flies from 3 biological replicates per timepoint. (**O**) Histogram of coefficients of diffusion for *LpWF*^*R3P51*^, n=500 tracks from 5 biological replicates.

### An evolutionary screen isolates mutants with loss of colonization

To identify the bacterial genetic basis for *LpWF* adhesion, we first sequenced *LpATCC8014* and *LpWF* using Illumina paired end 150 bp reads and performed de novo genome assembly. Bioinformatic comparisons between the genomes yielded 596 unique genetic differences, corresponding to 368 and 228 unique genes in *LpATCC8014* and *LpWF*, respectively. Most of the genetic differences were in uncharacterized genes (531 genes, 89%). 11% of the 596 genes were classified by RAST (*40*), and these corresponded to 13 metabolic pathways, including pathways involved in carbohydrate and nitrogen metabolism, DNA metabolism, and cell wall formation (**Table S1**). None of these provided a clear hypothesis regarding the genes responsible for colonization.

We next screened for colonization genes using an evolve and resequence approach (**Fig. 1L**). We reasoned that in the wild, *LpWF* requires the fly as a vector, imparting strong selection pressure to maintain colonization. We used continuous growth in a rich liquid medium to eliminate the selection pressure for attaching to the fly while maintaining selection for robust growth. After 51 passages, we observed a maintenance of growth rate (**Fig. S3, Table S2**) and a decrease in colonization efficiency for all 12 replicates (**Fig. 1M, Fig. S4, Table S3**). Replicate 3 at passage 51 (*LpWF*^*R3P51*^) was the weakest colonizer between biological replicates (**Fig. 1M, Fig. S4**). Over the course of the evolution experiment, *LpWF*^*R3*^ decreased in colonization efficiency beginning at passage 37 (**Fig. 1N, Fig. S4**), suggesting that a mutant with loss of function in one or more colonization genes took over the population at this time point. Bellymount imaging revealed a sharp decrease in gut adhesion and an increase in diffusion rate to 0.122 µm^2^/s for *LpWF*^*R3P51*^vs. 0.001µm^2^/s for *LpWF*) (**Fig. 1O, Movie S3**).

### Deep sequencing of mutants identifies a colonization island

To identify genomic changes associated with the colonization defects observed in the evolved populations, we resequenced the evolved strains and mapped the reads to the *LpWF* assembly, revealing only 11 single nucleotide polymorphisms (SNPs) with no obvious linkage to colonization (**Table S4**). The *LpWF* Illumina assembly was highly fragmented, with 316 contigs (**Table S5**). To search for structural variants, we resequenced both *LpWF* and *LpWF*^*R3P51*^ with PacBio HiFi long reads (**Table S5**), resulting in an assembly for *LpWF* composed of a single circular chromosome (3.23 Mbp), plus five plasmids (Plasmid 1: 185.5 Kbp, Plasmid 2: 60.0 Kbp, Plasmid 3: 41.6 Kbp, Plasmid 4: 17.2 Kbp, and Plasmid 5: 15.8 Kbp) (**Fig. 2A**). Analysis by PCR and read mapping to our reference genome for *LpWF* showed that Plasmid 1, which we named pKG-WF, is linear while the four remaining plasmids are circular (**Fig. S5, S6, Table S6**). Linear plasmids are known from several lactic acid bacteria, including *L. salivarus* (*41, 42*), *L. plantarum* (*43*), and *L. gasseri* (*44*) among others. In *LpWF*^*R3P51*^, the assembly revealed a loss of a contiguous 84.5 Kbp segment in pKG-WF (**Fig. 2A**). No short reads or long reads from *LpWF*^*R3P51*^ aligned to this region (**Fig. S7**). Alignments of short reads from passage 51 of each of the 12 replicates showed significant loss of the colonization island (**Fig. S7**). Loss of the island was detected by sequencing as early as the second week of passaging in R3P51, with a negative selection coefficient of 4% per generation (**Fig. S8**), consistent with the faster growth rate of the mutants (**Fig. S3**).

**Fig. 2.**
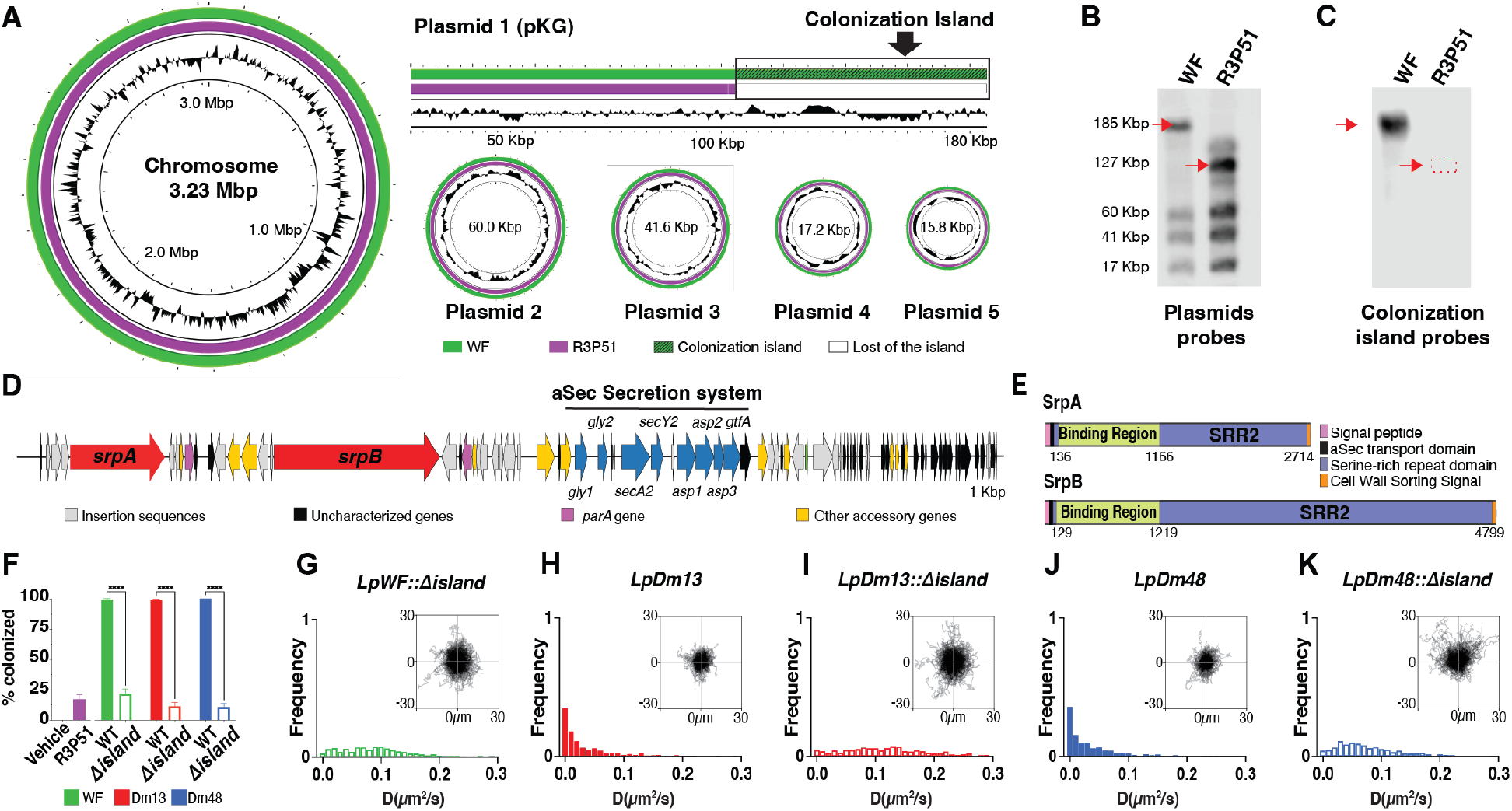
A colonization island containing serine-rich repeat proteins drives *L. plantarum* adhesion to the gut. (**A**) Long read sequencing of *LpWF* and the experimentally-evolved mutants indicates a colonization island. (**B**) PFGE and Southern blotting of plasmid partition genes shows a reduction in length of the linear pKG plasmid that contains the colonization island and (**C**) loss of the colonization island using probes for the island-specific genes *srpA* and *srpB*. (**D**) Map of the island with functional annotations. See Fig. S11B for additional annotations. (**E**) SRRP protein schematics. (**F**) Colonization is lost when the island is deleted in 3 separate *L. plantarum* isolates from wild *D. melanogaster*. Live fly, single bacterial cell diffusion coefficient histograms for (**G**) *LpWF::Δisland*, (**H**) *LpDm13*, (**I**) *LpDm13::Δisland*, (**J**) *LpDm48*, and (**K**) *LpDm48::Δisland*.

Loss of the putative colonization island from plasmid pKG-R3P51 was confirmed by Pulsed-Field Gel Electrophoresis (PFGE), which identified a reduction in the molecular weight of plasmid pKG (**Fig. 2B**). The size of the PFGE bands is consistent with the PacBio genome assembly, showing that the length of plasmid pKG-R3P51 is smaller than the plasmid pKG-WF (127.8 Kbp and 185.5 Kbp, respectively) (**Fig. S9**). Southern blot analysis confirmed that plasmid pKG-R3P51 is smaller in length than plasmid pKG-WF and that the colonization island is lost, confirming our bioinformatics analysis (**Fig. 2C**).

Functional annotation of the putative colonization island using InterProScan(*45*) revealed 80 open reading frames (**Fig. 2D**), including two putative large adhesins belonging to the group of Serine-Rich Repeat Proteins (SRRPs), which are known from *Streptococcus, Staphylococcus*, and one lactobacillus species, yet their specific role in host association remains little-understood (*23, 46, 47*). The two *LpWF* SRRPs, which we call *srpA* and *srpB*, are 2714 aa and 4799 aa respectively, together making up 26.7% of the island sequence, which is large for bacterial proteins but normal for SRRPs (**Fig. 2E**). The island also contained the auxiliary SRRP secretion genes called the aSec system (**Fig. 2D;** (*22, 23, 48*–*50*)). The domain organization of *srpA* and *srpB* includes a signal sequence peptide at the N-terminus, an accessory Sec transport domain (AST), two serine-rich repeat domains (SRR-1 and SRR-2, respectively), a binding region domain (BR), and an LPXTG cell wall anchoring domain (**Fig. 2E**).

### The colonization island is necessary for colonization

To determine the role that deletion of the island played in the *LpWF*^*R3P51*^ colonization phenotypes versus the role of the 11 point mutants, we developed a Cas9-based approach to remove the complete island by truncating the pKG plasmid in the *LpWF* background (**Fig. S10**). This mutant, *LpWF::Δisland*, exhibits a ∼85% reduction in colonization efficiency relative to wild-type, indistinguishable from the *LpWF*^*R3P51*^ mutant, indicating that the island is necessary for strong colonization (**Fig. 2F**). Bellymount of *LpWF::Δisland* confirmed that the island is necessary for stable attachment to the crop luminal surface (median diffusion coefficient 0.073 µm^2^/s vs. 0.0010 µm^2^/s for *LpWF*), (**Fig. 2G, Movie S4**).

Instability of the island during 51 days of evolution appeared to arise from many repeated insertion sequences (ISs), which are small transposable elements, classified into six different families (IS30, IS1595, IS982, ISL3, IS5, and IS1182). We predicted recombination sites in the island and visualized them on a dot plot (**Fig. S11**). Based on the position of the ISs, we divided the colonization island into 13 different blocks of genes flanked by ISs (I-XIII), yielding 16 putative recombination sites, most of them situated between blocks II to VIII (**Fig. S11**).

To identify mutants with partial deletions of the island, we used qPCR directed at specific IS blocks to screen the early timepoints in the evolution experiment when the populations were heterogeneous (**Fig. S12; Table S7**). We isolated two main genotypes, which we further confirmed using long-read sequencing (**Fig. S13**): i) mutant D11 had deletions of blocks III – XIII, which includes *secA2* and carries the same colonization defects as *LpWF::Δisland*, indicating *secA2* is necessary for function of the island (**Fig. S14, Movie S5**). ii) Mutant F9 had deletions of blocks II - IV, including *srpB*. F9 retained *srpA* and the *secA2* secretion machinery. Accordingly, F9 colonized a greater proportion of flies than *LpWF::Δisland* or D11, indicating that one SRRP and the aSec system are sufficient for colonization. F9 exhibited spatial specificity for the crop but not the proventriculus (**Fig S14, Movie S6**), suggesting spatial specificity of the two SRRPs.

### The colonization island is necessary for colonization specificity in other *L. plantarum* strains

The colonization island is not present in short read genomes for the technical reason that extensive sequence repeats in the SRRPs and ISs cause fragmented assemblies with very short contigs. To directly assess the prevalence of the colonization island in other *L. plantarum* strains without assembly, we downloaded the raw reads for NCBI genomes from SRA and aligned them to the *LpWF* island. We first constructed a WGS database with 1,247 raw read data sets sequenced by Illumina (96.1%), PacBio (2.4%), Oxford Nanopore (0.89%), and other long read technologies (0.61%) (**Table S8**). Next, we mapped the reads against the nucleotide sequence of the island. We identified two *L. plantarum* strains, named *LpDm13* and *LpDm48*, which were isolated from wild *D. melanogaste*r flies collected in Ithaca, NY and generously shared by the authors (McMullen et al., 2021) (**Fig. S15**). We confirmed the presence of the colonization islands by long-read sequencing. The *LpDm13* and *LpDm48* islands are each a comparable size to *LpWF’s* island (97.3 Kbp and 113.8 Kbp, respectively) and within a linear plasmid (**Fig. S16**). The island in *LpDm13* consists of 79 ORFs, while *LpDm48’s* island comprises 108 ORFs. Both islands harbor two SRRP alleles (SrpA= 513 kD and SrpB= 442 kD for *LpDm13*; and SrpA=352 kD and SrpB= 493 kD for *LpDm48*) and a corresponding aSec secretion system, with numerous adjacent ISs.

*LpDm13* and *LpDm48* both colonized nearly 100% of flies when inoculated at a low dose (**Fig. 2F**) with stable populations averaging ∼20,000-50,000 CFUs inside the fly gut, consistent with *LpWF* (**Fig. S17**), and showed similar spatial localization (**Fig. S18**) and diffusion (**Fig. 2H, 2J, Movies S7, S8**). We next constructed mutants in *LpDm13* and *LpDm48* with a knockout of the complete island. These mutants exhibited an acute colonization deficiency (**Fig. 2F; S17, S18**) and adhesion deficiency by Bellymount (**Fig. 2I, 2K, Movies S9, S10**) compared to the corresponding wildtype strains. Although the *LpDm48*::*Δisland* mutant lost its capacity to colonize, it retained its ability to form aggregates, indicating the aggregation trait is not encoded on the island, as suggested for some other SRRP studies (*46*). Overall, our results indicate a conserved colonization specificity island within *L. plantarum*.

### The colonization island is conserved across the Firmicutes

SRRPs with aSec systems were previously known from just three genera, all within the Lactobacillales (*46*). To explore conservation of the colonization island across other bacterial genera, we first used BLASTP to search for the complete aSec system in the Pathosystems Resource Integration Center (PATRIC) database of bacterial genomes, focused on human bacteria (*51*). Most bacterial genome assemblies were highly fragmented due to repeat elements, which were also abundant in the *LpWF* colonization island. As a conservative approach to identify homologous colonization islands, we excluded BLAST hits where the aSec genes were not in a semicontiguous block because some aSec genes have homologs in the genome (*23*). We first considered cases with at least one complete SRRP gene with the aSec block. Due to the lack of amino acid sequence conservation in SRRP genes, we used a hidden Markov model based search to detect these (Supplemental Methods). Despite the preponderance of highly fragmented genome assemblies, we discovered the complete colonization island in 207 bacterial genomes spanning the entire Firmicutes phylum, including 134 Lactobacillales and 25 Clostridia, as well as in three Actinomycetes and one *ε*-Proteobacterium (**Fig 3A, S19, Table S9**). We found the aSec system with a non-SRRP, large adhesin in an additional 42 genomes, including 7 Clostridia and 1 *γ*-Proteobacterium (**Fig. S19**). While there was broad synteny of the aSec genes across the phylogeny (**Fig. 3A**), there were stark differences in the adhesins, consistent with these being host-specific.

**Fig. 3.**
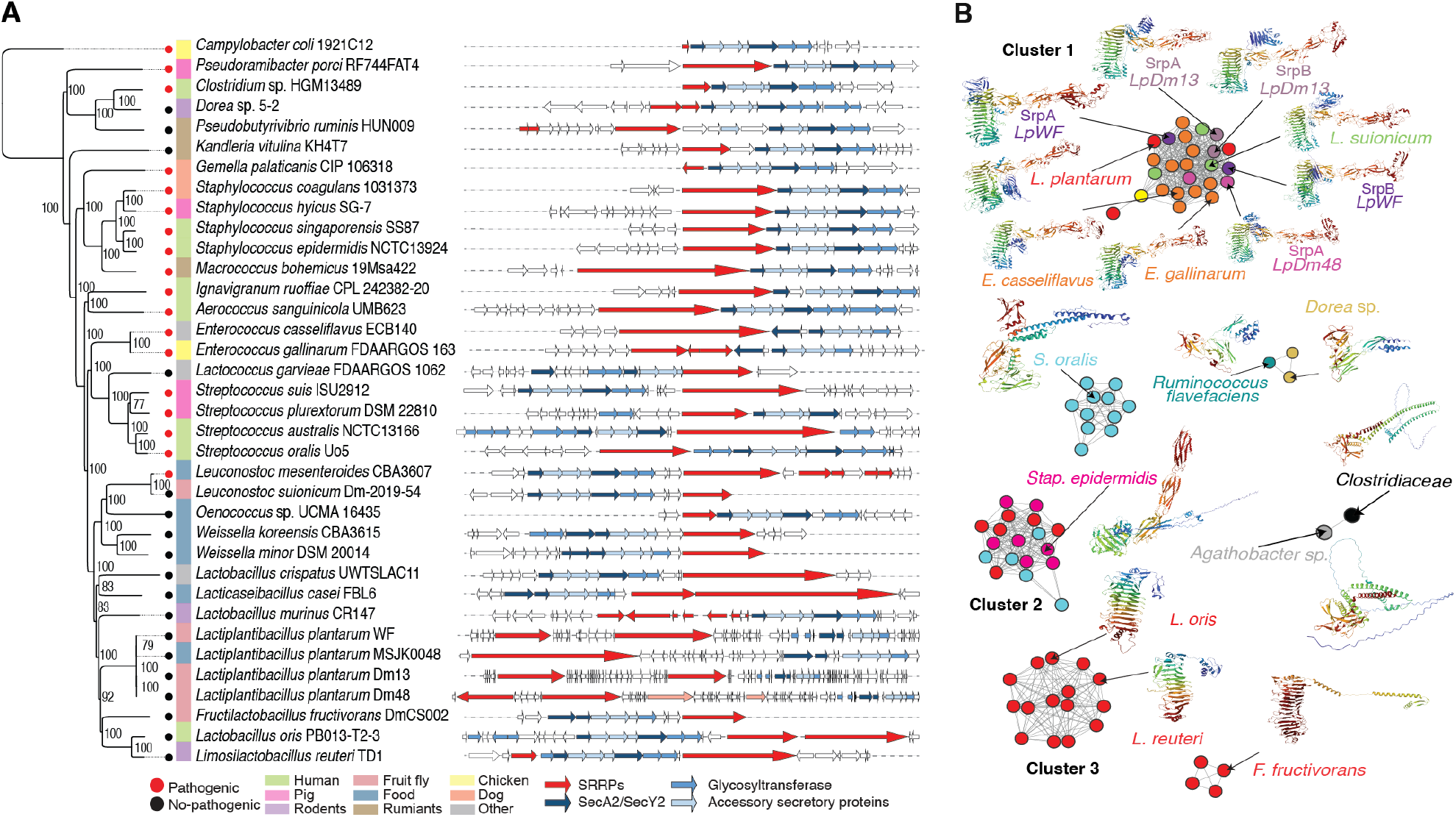
The colonization island is widely conserved in host-associated bacteria. (**A**) Core genome phylogeny of 36 species and maps of their corresponding colonization islands. (**B**) Clustering analysis of the binding regions of diverse bacterial colonization islands and representative predicted structures for each of the clusters. See Fig. S20 for complete results.

To investigate the conservation of the SRRPs, we constructed a similarity network based on structural motifs in the binding regions, revealing 15 clusters and 2 singletons (**Fig. 3B, S20**). We simulated their structures using RosettaFold. Three of these clusters, including the *LpWF* cluster, contain a beta-solenoid motif, which was present in genera including *Enterococcus, Fructilactibacillus, Lactobacillus, Lactoplantibacillus, Leuconostoc* and *Weisella*. Another structural motif was a chain of immunoglobulin-like domains, which was present in genera including *Oenococcus, Pediococcus*, and *Staphylococcus*, as well as in combination with the beta-solenoid domain in the *LpWF* cluster. Additional unique binding regions were present in the Clostridiales, including in *Dorea* and *Pseudoramibacter*. We analyzed the conservation of amino acids within each of the three largest clusters of binding regions (Clusters 1 – 3 in **Fig. 3B**) and found a strong signature of positive selection (**Fig. S21**), consistent with the role of these proteins in host specificity. Non-synonymous nucleotide changes were concentrated in the beta-solenoid motifs of Clusters 1 and 3 and in the first immunoglobulin-like domain of Cluster 2 (**Fig. S21**).

Based on the conservation of this genomic region, we defined the minimum colonization island gene set, composed of seven core genes with at least one *srp*, the tandem transporters *secA2*/*secY2*, the glycosyltransferase *gtfA*, and three accessory secretory protein genes: *asp1, asp2*, and *asp3*. The order of the genes was conserved at the genus level, but differed between some families, suggesting rearrangements within the island on longer evolutionary timescales (**Fig. 3A**).

To assess whether the island is predominantly passed between strains by vertical transfer or horizontal gene transfer, we performed co-phylogenetic analysis. Vertical transfer would produce parallel phylogenies for the core genome and the colonization island genes whereas horizontal transfers of the island would produce discrepancies (*52*). We first reconstructed the phylogeny of the host species using the core genome, which was composed of 81 orthologous proteins (**Table S10**), and the island using the six conserved aSec proteins (**Fig. S22**). We then compared the two phylogenies for each well-resolved clade using a cutoff bootstrap value of > 70, and estimated the Patristic distance (*53, 54*). The branching pattern of the aSec phylogeny overall matched the genome phylogeny (**Fig. S22**; mean ParaFit Global index: 0.021, p-value < 0.05), indicating that the colonization island was likely present in the ancestor of the Firmicutes, a largely host-associated phylum. Some cases of horizontal gene transfer were apparent, such as the acquisition of the island by the gram negative *Campylobacter coli* from the gram positive *Staphylococcus epidermidis*, indicating spread of the island beyond the Firmicutes.

## Conclusions

Our study identified a mechanism of symbiont gut colonization through specific binding of a physical niche tissue, deep within the gut. The ability to directly observe the colonization process at the single bacterial cell level allows a clear differentiation between a dynamically stable population that transiently binds the gut wall and maintains itself by growing faster than it is shed versus a population that is statically adhered to the gut wall. The genes for static attachment that we identified have likely gone undiscovered across the Firmicutes for two primary reasons. First, significant loss of the colonization genes occurred in just two weeks of culture in rich media (**Fig. S8)**, indicating strong selection against the colonization genes when grown in lab conditions outside the natural host. The slower growth of the wildtype strain versus the mutants (**Fig. S3**) may stem from the large size of the SRRPs (nearly 500 kD for SrpB) and considerable energy required to produce them. Second, the serine-rich repeats and other repetitive sequences cause fragmented genome assemblies when sequenced with the predominant short-read sequencing technologies. The study of wild strains using current long read technologies can overcome these two obstacles.

The genes for this specificity include the aSec system, which was known for secretion of virulence factors in *Streptococcus* and *Staphylococcus* species that are predominantly opportunistic pathogens (*23*). The evolutionary selection for molecules that lead opportunistic pathogens to cross an epithelial barrier and cause a systemic infection are different from the long term selective pressures that maintain them as non-pathogenic commensals because the opportunistic infections are typically not transmitted as such (*17*). Rather, they arise anew in each case, for instance in immunocompromised individuals. Such opportunistic infections therefore lack long term evolutionary pressures to refine their targeting of e.g. host blood cells (*16*). Symbiotic bacteria on the other hand must reach a host niche without becoming stuck in low quality habitat elsewhere in the digestive tract. To colonize a new host, a symbiont is thus selected for its ability to selectively colonize the symbiotic niche and not stick to other lower quality sites in the host (*11, 55*)

For commensals, receptor-ligand binding interactions provide a mechanism that hones specificity, which can help commensals locate their correct niche without becoming entrapped in undesirable locations in the rest of the gut, allowing the establishment of colonization with very few initial inoculants (*37*). The conservation of these genes into the Clostridia and even Proteobacteria suggests a mechanism by which the observed patterns in strain engraftment may occur (*56*).

## Acknowledgments

Prof. John Chaston (Brigham Young University) provided the *LpDm13* and *LpDm48* strains isolated from wild *D. melanogaster*. Prof. Reingard Grabherr and Prof. Stefan Heinl (BOKU Vienna) provided the pCD256-mCherry plasmid.

## Funding

National Institutes of Health grant R01 (WBL)

National Institutes of Health grant R21 (WBL)

National Science Foundation grant IOS (WBL)

National Science Foundation CAREER Award grant IOS (WBL)

The Carnegie Institution for Science Endowment

National Institutes of Health T32 training grant to JHU CMDB program

## Author contributions

Conceptualization: KGG, KA, WBL

Investigation: KGG, KA, RD, BO, AD, SA, XY, RW

Supervision: WBL, NG

Writing: KGG, KA, RD, WBL

## Data and materials availability

All data and code are available from KGG’s GitHub repository (https://github.com/K-Gutierrez/LpColonizationIsland/). The BioProject accession number PRJNA1094941, corresponds to the raw data from all the evolved replicates and passages of the evolution experiment. The BioProject accession number PRJNA1094948, corresponds to the *LpWF* and *LpWF*^*R3P51*^ short read assemblies. The BioProject accession number PRJNA1093452, corresponds to the *LpWF, LpWF*^*R3P51*^, *LpDm13, LpDm48, LpWF-D11*, and *LpWF-F9* long read assemblies. All strains are available upon request.

## Supplementary Materials

Materials and Methods

Figs. S1 to S22

Tables S1 to S15

References (*1-44*)

Movies S1 to S10

Data S1

## References and Notes

1. W. Burroughs, N. A. Frank, P. Gerlaugh, R. M. Bethke, Preliminary observations upon factors influencing cellulose digestion by rumen microorganisms. J. Nutr. 40, 9–24 (1950).

2. W. H. Karasov, C. Martínez Del Rio, E. Caviedes-Vidal, Ecological physiology of diet and digestive systems. Annu. Rev. Physiol. 73, 69–93 (2011).

3. O. M. Sokolovskaya, A. N. Shelton, M. E. Taga, Sharing vitamins: Cobamides unveil microbial interactions. Science (80-.). 369 (2020), doi:10.1126/science.aba0165.

4. A. J. Mcdermott, G. B. Huffnagle, The microbiome and regulation of mucosal immunity. Immunology. 142, 24–31 (2014).

5. J. Walter, M. X. Maldonado-Gómez, I. Martínez, To engraft or not to engraft: an ecological framework for gut microbiome modulation with live microbes. Curr. Opin. Biotechnol. 49, 129–139 (2018).

6. M. Wolter, E. T. Grant, M. Boudaud, A. Steimle, G. V. Pereira, E. C. Martens, M. S. Desai, Leveraging diet to engineer the gut microbiome. Nat. Rev. Gastroenterol. Hepatol. 18, 885–902 (2021).

7. J. Walter, R. Ley, The Human Gut Microbiome: Ecology and Recent Evolutionary Changes. Annu. Rev. Microbiol. 65, 411–429 (2011).

8. P. J. Turnbaugh, V. K. Ridaura, J. J. Faith, F. E. Rey, R. Knight, J. I. Gordon, The effect of diet on the human gut microbiome: a metagenomic analysis in humanized gnotobiotic mice. Sci. Transl. Med. 1, 6ra14 (2009).

9. G. P. Donaldson, S. M. Lee, S. K. Mazmanian, Gut biogeography of the bacterial microbiota, 1–13 (2015).

10. G. McCallum, C. Tropini, The gut microbiota and its biogeography. Nat. Rev. Microbiol. 22, 105–118 (2023).

11. K. Takeshita, Y. Kikuchi, Riptortus pedestris and Burkholderia symbiont: an ideal model system for insect-microbe symbiotic associations. Res. Microbiol. 168, 175–187 (2017).

12. R. Dodge, E. W. Jones, H. Zhu, B. Obadia, D. J. Martinez, C. Wang, A. Aranda-Díaz, K. Aumiller, Z. Liu, M. Voltolini, E. L. Brodie, K. C. Huang, J. M. Carlson, D. A. Sivak, A. C. Spradling, W. B. Ludington, Nat. Commun., in press, doi:10.1038/s41467-023-36942-x.

13. C. Fung, S. Tan, M. Nakajima, E. C. Skoog, L. F. Camarillo-Guerrero, J. A. Klein, T. D. Lawley, J. V. Solnick, T. Fukami, M. R. Amieva, High-resolution mapping reveals that microniches in the gastric glands control Helicobacter pylori colonization of the stomach. PLoS Biol. 17, 1–28 (2019).

14. S. V. Nyholm, E. V. Stabb, E. G. Ruby, M. J. McFall-Ngai, Establishment of an animal-bacterial association: Recruiting symbiotic vibrios from the environment. Proc. Natl. Acad. Sci. U. S. A. 97, 10231–10235 (2000).

15. K. A. Kline, S. Fälker, S. Dahlberg, S. Normark, B. Henriques-Normark, Bacterial Adhesins in Host-Microbe Interactions. Cell Host Microbe. 5, 580–592 (2009).

16. E. H. Beachey, Bacterial adherence: adhesin-receptor interactions mediating the attachment of bacteria to mucosal surface. J. Infect. Dis. 143, 325–345 (1981).

17. S. P. Brown, D. M. Cornforth, N. Mideo, Evolution of virulence in opportunistic pathogens: Generalism, plasticity, and control. Trends Microbiol. 20, 336–342 (2012).

18. J. G. Sanders, D. D. Sprockett, Y. Li, D. Mjungu, E. V Lonsdorf, J. N. Ndjango, A. V Georgiev, J. A. Hart, C. M. Sanz, D. B. Morgan, M. Peeters, B. H. Hahn, A. H. Moeller, Widespread extinctions of co-diversified primate gut bacterial symbionts from humans. Nat. Microbiol., 1039–1050 (2023).

19. T. A. Suzuki, J. L. Fitzstevens, V. T. Schmidt, H. Enav, K. E. Huus, M. M. Ngwese, A. Grießhammer, A. Pfleiderer, B. R. Adegbite, J. F. Zinsou, M. Esen, T. P. Velavan, A. A. Adegnika, L. H. Song, T. D. Spector, A. L. Muehlbauer, N. Marchi, H. Kang, L. Maier, R. Blekhman, L. Ségurel, G. P. Ko, N. D. Youngblut, P. Kremsner, R. E. Ley, Codiversification of gut microbiota with humans. Science (80-.). 377, 1328–1332 (2022).

20. M. K. Nishiguchi, E. G. Ruby, M. J. McFall-Ngai, Competitive Dominance among Strains of Luminous Bacteria Provides an Unusual Form of Evidence for Parallel Evolution in Sepiolid Squid-Vibrio Symbioses. 64, 3209–3213 (1998).

21. M. Juhas, J. R. Van Der Meer, M. Gaillard, R. M. Harding, D. W. Hood, D. W. Crook, Genomic islands: Tools of bacterial horizontal gene transfer and evolution. FEMS Microbiol. Rev. 33, 376–393 (2009).

22. C. J. Sanchez, P. Shivshankar, K. Stol, S. Trakhtenbroit, P. M. Sullam, K. Sauer, P. W. M. Hermans, C. J. Orihuela, The pneumococcal serine-rich repeat protein is an intraspecies bacterial adhesin that promotes bacterial aggregation in Vivo and in biofilms. PLoS Pathog. 6, 33–34 (2010).

23. M. Braunstein, B. A. Bensing, P. M. Sullam, The Two Distinct Types of SecA2-Dependent Export Systems. Microbiol. Spectr. 7 (2019), doi:10.1128/microbiolspec.psib-0025-2018.

24. H. Ochman, N. A. Moran, Genes lost and genes found: Evolution of bacterial pathogenesis and symbiosis. Science (80-.). 292, 1096–1098 (2001).

25. E. G. Ruby, M. Urbanowski, J. Campbell, A. Dunn, M. Faini, R. Gunsalus, P. Lostroh, C. Lupp, J. McCann, D. Millikan, A. Schaefer, E. Stabb, A. Stevens, K. Visick, C. Whistler, E. P. Greenberg, Complete genome sequence of Vibrio fischeri: A symbiotic bacterium with pathogenic congeners. Proc. Natl. Acad. Sci. U. S. A. 102, 3004–3009 (2005).

26. L. Speare, A. G. Cecere, K. R. Guckes, S. Smith, M. S. Wollenberg, M. J. Mandel, T. Miyashiro, A. N. Septer, Bacterial symbionts use a type VI secretion system to eliminate competitors in their natural host. Proc. Natl. Acad. Sci. U. S. A. 115, E8528–E8537 (2018).

27. P. J. Planet, S. C. Kachlany, D. H. Fine, R. DeSalle, D. H. Figurski, The widespread colonization island of Actinobacillus actinomycetemcomitans. Nat. Genet. 34, 193–198 (2003).

28. T. J. Wiles, M. Jemielita, R. P. Baker, B. H. Schlomann, S. L. Logan, J. Ganz, E. Melancon, J. S. Eisen, K. Guillemin, R. Parthasarathy, Host Gut Motility Promotes Competitive Exclusion within a Model Intestinal Microbiota. PLoS Biol. 14, e1002517 (2016).

29. B. H. Schlomann, T. J. Wiles, E. S. Wall, K. Guillemin, R. Parthasarathy, Bacterial Cohesion Predicts Spatial Distribution in the Larval Zebrafish Intestine. Biophys. J. 115, 2271–2277 (2018).

30. N. A. Broderick, B. Lemaitre, Gut-associated microbes of Drosophila melanogaster. Gut Microbes. 3, 307–321 (2012).

31. H. A. Seddik, F. Bendali, F. Gancel, I. Fliss, G. Spano, D. Drider, Lactobacillus plantarum and Its Probiotic and Food Potentialities, 1–12 (2017).

32. M. Schwarzer, K. Makki, G. Storelli, I. Machuca-Gayet, D. Srutkova, P. Hermanova, M. E. Martino, S. Balmand, T. Hudcovic, A. Heddi, J. Rieusset, H. Kozakova, H. Vidal, F. Leulier, Lactobacillus plantarum strain maintains growth of infant mice during chronic undernutrition. Science (80-.). 351, 854–857 (2016).

33. M. E. Martino, P. Joncour, R. Leenay, H. Gervais, M. Shah, S. Hughes, B. Gillet, C. Beisel, F. Leulier, Bacterial Adaptation to the Host’s Diet Is a Key Evolutionary Force Shaping Drosophila-Lactobacillus Symbiosis. Cell Host Microbe. 24, 109-119.e6 (2018).

34. J.-H. Ryu, K.-B. Nam, C.-T. Oh, H.-J. Nam, S.-H. Kim, J.-H. Yoon, J.-K. Seong, M.-A. Yoo, I.-H. Jang, P. T. Brey, W.-J. Lee, The homeobox gene Caudal regulates constitutive local expression of antimicrobial peptide genes in Drosophila epithelia. Mol. Cell. Biol. 24, 172–185 (2004).

35. G. Storelli, M. Strigini, T. Grenier, L. Bozonnet, M. Schwarzer, C. Daniel, R. Matos, F. Leulier, Drosophila Perpetuates Nutritional Mutualism by Promoting the Fitness of Its Intestinal Symbiont Lactobacillus plantarum. Cell Metab. 27, 1–16 (2018).

36. M. E. Martino, J. R. Bayjanov, B. E. Caffrey, M. Wels, P. Joncour, S. Hughes, B. Gillet, M. Kleerebezem, S. A. F. T. van Hijum, F. Leulier, Nomadic lifestyle of Lactobacillus plantarum revealed by comparative genomics of 54 strains isolated from different habitats. Environ. Microbiol. 18, 4974–4989 (2016).

37. B. Obadia, Z. T. Güvener, V. Zhang, J. A. Ceja-Navarro, E. L. Brodie, W. W. Ja, W. B. Ludington, Probabilistic Invasion Underlies Natural Gut Microbiome Stability. Curr. Biol. 27 (2017), doi:10.1016/j.cub.2017.05.034.

38. L. A. J. Koyama, A. Aranda-Díaz, Y.-H. Su, S. Balachandra, J. L. Martin, W. B. Ludington, K. C. Huang, L. E. O’Brien, Bellymount enables longitudinal, intravital imaging of abdominal organs and the gut microbiota in adult Drosophila. PLoS Biol. 18, e3000567–19 (2020).

39. N. Buntin, W. M. de Vos, T. Hongpattarakere, Variation of mucin adhesion, cell surface characteristics, and molecular mechanisms among Lactobacillus plantarum isolated from different habitats. Appl. Microbiol. Biotechnol. 101, 7663–7674 (2017).

40. A. Wilke, J. Bischof, W. Gerlach, E. Glass, T. Harrison, K. P. Keegan, T. Paczian, W. L. Trimble, S. Bagchi, A. Grama, S. Chaterji, F. Meyer, The MG-RAST metagenomics database and portal in 2015. Nucleic Acids Res. 44, D590–D594 (2016).

41. Y. Li, C. Canchaya, F. Fang, E. Raftis, K. A. Ryan, J. P. Van Pijkeren, D. Van Sinderen, P. W. O’Toole, Distribution of megaplasmids in Lactobacillus salivarius and other Lactobacilli. J. Bacteriol. 189, 6128–6139 (2007).

42. E. J. Raftis, B. M. Forde, M. J. Claesson, P. W. O’Toole, Unusual genome complexity in Lactobacillus salivarius JCM1046. BMC Genomics. 15, 1–15 (2014).

43. D. Davray, H. Bawane, R. Kulkarni, Non-redundant nature of Lactiplantibacillus plantarum plasmidome revealed by comparative genomic analysis of 105 strains. Food Microbiol. 109, 104153 (2023).

44. Y. G. Abs El-Osta, A. J. Hillier, B. E. Davidson, M. Dobos, Pulsed-field gel electrophoretic analysis of the genome of Lactobacillus gasseri ATCC33323, and construction of a physical map. Electrophoresis. 23, 3321–3331 (2002).

45. M. Blum, H. Y. Chang, S. Chuguransky, T. Grego, S. Kandasaamy, A. Mitchell, G. Nuka, T. Paysan-Lafosse, M. Qureshi, S. Raj, L. Richardson, G. A. Salazar, L. Williams, P. Bork, A. Bridge, J. Gough, D. H. Haft, I. Letunic, A. Marchler-Bauer, H. Mi, D. A. Natale, M. Necci, C. A. Orengo, A. P. Pandurangan, C. Rivoire, C. J. A. Sigrist, I. Sillitoe, N. Thanki, P. D. Thomas, S. C. E. Tosatto, C. H. Wu, A. Bateman, R. D. Finn, The InterPro protein families and domains database: 20 years on. Nucleic Acids Res. 49, D344–D354 (2021).

46. M. S. Cinar, A. Niyas, F. Y. Avci, Serine-rich repeat proteins: well-known yet little-understood bacterial adhesins. J. Bacteriol., 1–15 (2023).

47. D. Latousakis, D. A. MacKenzie, A. Telatin, N. Juge, Serine-rich repeat proteins from gut microbes. Gut Microbes. 11, 102–117 (2020).

48. S. Sequeira, D. Kavanaugh, D. A. MacKenzie, T. Šuligoj, S. Walpole, C. Leclaire, A. P. Gunning, D. Latousakis, W. G. T. Willats, J. Angulo, C. Dong, N. Juge, Structural basis for the role of serine-rich repeat proteins from Lactobacillus reuteri in gut microbe-host interactions. Proc. Natl. Acad. Sci. U. S. A. 115, E2706–E2715 (2018).

49. A. K. Singh, S. A. Woodiga, M. A. Grau, S. J. Kinga, Streptococcus oralis neuraminidase modulates adherence to multiple carbohydrates on platelets. Infect. Immun. 85, 1–15 (2017).

50. B. A. Bensing, L. V. Loukachevitch, K. M. Mcculloch, H. Yu, K. R. Vann, Z. Wawrzak, S. Anderson, X. Chen, P. M. Sullam, T. M. Iverson, Structural basis for sialoglycan binding by the streptococcus sanguinis SrpA adhesin. J. Biol. Chem. 291, 7230–7240 (2016).

51. J. J. Gillespie, A. R. Wattam, S. A. Cammer, J. L. Gabbard, M. P. Shukla, O. Dalay, T. Driscoll, D. Hix, S. P. Mane, C. Mao, E. K. Nordberg, M. Scott, J. R. Schulman, E. E. Snyder, D. E. Sullivan, C. Wang, A. Warren, K. P. Williams, T. Xue, H. S. Yoo, C. Zhang, Y. Zhang, R. Will, R. W. Kenyon, B. W. Sobral, Patric: The comprehensive bacterial bioinformatics resource with a focus on human pathogenic species. Infect. Immun. 79, 4286–4298 (2011).

52. S. M. Soucy, J. Huang, J. P. Gogarten, Horizontal gene transfer: Building the web of life. Nat. Rev. Genet. 16, 472–482 (2015).

53. P. Legendre, Y. Desdevises, E. Bazin, A statistical test for host-parasite coevolution. Syst. Biol. 51, 217–234 (2002).

54. E. Paradis, Analysis of phylogenetics and evolution with R (Springer, New York, Second edi., 2012).

55. S. V. Nyholm, M. J. McFall-Ngai, A lasting symbiosis: how the Hawaiian bobtail squid finds and keeps its bioluminescent bacterial partner. Nat. Rev. Microbiol. 19, 666–679 (2021).

56. V. Aggarwala, I. Mogno, Z. Li, C. Yang, G. J. Britton, A. Chen-Liaw, J. Mitcham, G. Bongers, D. Gevers, J. C. Clemente, J. F. Colombel, A. Grinspan, J. Faith, Precise quantification of bacterial strains after fecal microbiota transplantation delineates long-term engraftment and explains outcomes. Nat. Microbiol. 6, 1309–1318 (2021).

